# Automated Deep Learning-Based Diagnosis and Molecular Characterization of Acute Myeloid Leukemia using Flow Cytometry

**DOI:** 10.1101/2023.09.18.558289

**Authors:** Joshua E. Lewis, Lee A.D. Cooper, David L. Jaye, Olga Pozdnyakova

## Abstract

Current flow cytometric analysis of blood and bone marrow samples for diagnosis of acute myeloid leukemia (AML) relies heavily on manual intervention in both the processing and analysis steps, introducing significant subjectivity into resulting diagnoses and necessitating highly trained personnel. Furthermore, concurrent molecular characterization via cytogenetics and targeted sequencing can take multiple days, delaying patient diagnosis and treatment. Attention-based multi-instance learning models (ABMILMs) are deep learning models which make accurate predictions and generate interpretable insights regarding the classification of a sample from individual events/cells; nonetheless, these models have yet to be applied to flow cytometry data. In this study, we developed a computational pipeline using ABMILMs for the automated diagnosis of AML cases based exclusively on flow cytometric data. Analysis of 1,820 flow cytometry samples shows that this pipeline provides accurate diagnoses of acute leukemia [AUROC 0.961] and accurately differentiates AML *versus* B- and T-lymphoblastic leukemia [AUROC 0.965]. Models for prediction of 9 cytogenetic aberrancies and 32 pathogenic variants in AML provide accurate predictions, particularly for t(15;17)(*PML*::*RARA*) [AUROC 0.929], t(8;21)(*RUNX1*::*RUNX1T1*) [AUROC 0.814], and *NPM1* variants [AUROC 0.807]. Finally, we demonstrate how these models generate interpretable insights into which individual flow cytometric events and markers deliver optimal diagnostic utility, providing hematopathologists with a data visualization tool for improved data interpretation, as well as novel biological associations between flow cytometric marker expression and cytogenetic/molecular variants in AML. Our study is the first to illustrate the feasibility of using deep learning-based analysis of flow cytometric data for automated AML diagnosis and molecular characterization.

## INTRODUCTION

The diagnosis of acute myeloid leukemia (AML) requires integration of myeloblast quantification and phenotypic characterization via flow cytometry and microscopic assessment, as well as genomic characterization via cytogenetics and targeted sequencing panels^1^. Flow cytometry allows for rapid measurement of physical properties and cell surface marker expression of hundreds of thousands of hematopoietic cells from blood and bone marrow aspirate specimens within a few hours^2^. This permits detection and characterization of aberrant myeloblast populations suggestive of AML on a time frame that provides clinicians the ability to expeditiously alter patient management. On the other hand, molecular characterization for detecting the presence of cytogenetic aberrancies and/or pathogenic variants in myeloblast populations often takes multiple days. This delay particularly complicates management of AML patients with certain genetic abnormalities requiring targeted therapies and/or changes in standard induction chemotherapy regimens^1,3^.

Current flow cytometric analysis relies heavily on manual intervention in both data processing, including compensation and gating, and data interpretation; this limits analysis of flow cytometry data to highly trained laboratory technologists and hematopathologists. Having an automated approach for analysis of flow cytometry data which does not rely on manual intervention would expand its use to a wider group of laboratorians and diagnosticians, similar to what automated pre-classification of white blood cell differentials has done for peripheral blood smears^4^. Additionally, manual data interpretation by hematopathologists introduces an element of subjectivity which could impact patient diagnosis and treatment; automated interpretation of data would eliminate such subjectivity. Finally, such automated software could be used as a diagnostic tool for improved data visualization and integration for hematopathologists, especially for complex cases.

Machine learning models have been extensively developed for applications in hematopathology, including for analysis of flow cytometry data^4–6^. Nonetheless, newer deep learning-based models, which have demonstrated improved predictive performance and interpretability compared to conventional machine learning techniques, have mainly been applied to morphologic assessment of blood and bone marrow aspirate samples rather than flow cytometric analysis^5^. A handful of studies have recently applied deep learning approaches to flow and mass cytometry data, including using convolutional neural networks (CNNs) on data histograms for detection of classic Hodgkin lymphoma^7^ and on CyTOF data for diagnosis of latent CMV infection^8^, combining self-organizing maps and CNNs for classification of mature B-cell neoplasms^9^, and using deep neural networks for minimal/measurable residual disease (MRD) analysis in acute lymphoblastic leukemia (ALL)^10^ and chronic lymphocytic leukemia^11^. However, attention-based multi-instance learning models (ABMILMs), a deep learning approach used to identify individual sample components which provide the most diagnostic value for individual cases (i.e., which flow cytometry events and markers are most important for the diagnosis of AML for a particular patient), have yet to be applied to flow cytometry data despite significant success in histopathology applications^12,13^.

In this study, we utilize ABMILMs for automated diagnosis and molecular characterization of AML cases from flow cytometry performed on blood and bone marrow samples. We developed a computational pipeline consisting of a series of deep learning models for predicting the presence or absence of acute leukemia, differentiating between AML and ALL cases, and finally predicting the presence or absence of 9 cytogenetic aberrancies and 32 pathogenic variants in AML cases. Pipeline implementation on 1,820 flow cytometry samples shows that these models provide strong predictive performance overall, as well as for challenging cases. Additionally, analysis of model-outputted attention values for individual flow cytometry events and their correlations with cell surface marker expression identifies which markers provide optimal utility for diagnosis and molecular characterization of AML cases. Finally, we give a representative example of a model-generated flow cytometry report providing model predictions and analysis tools for hematopathologists to improve flow cytometry data interpretation. Together, our results demonstrate the feasibility of using deep learning to make automated diagnoses of AML and provide insight into clinically actionable molecular variants based solely on flow cytometry data.

## MATERIALS AND METHODS

### Flow cytometry samples

Flow cytometry was performed on blood and bone marrow specimens in the hematology laboratory at Brigham and Women’s Hospital from 2019-2022 for routine patient care. Two-tube 10-color leukemia panels were run using the BD Biosciences (Franklin Lakes, NJ, USA) FACSCanto™ II Flow Cytometry System (Becton Dickinson) (**Supplementary Table 1)**. 1,820 flow cytometry cases were analyzed, 732 with definitive absence of acute leukemia, and 736 with definitive presence of acute leukemia (**Supplementary Table 2**). Among acute leukemia cases, 568 were diagnosed as acute myeloid leukemia (AML), and 168 were diagnosed as B- or T-lymphoblastic leukemia (ALL). The dataset was composed of initial diagnoses as well as follow-up cases, including pre- and post-treatment. A 20% blast count threshold based on flow cytometric analysis was used for all acute leukemia cases regardless of the presence of AML-defining translocations as well as for follow-up cases; positive AML cases with defining translocations or follow-up leukemia cases with less than 20% blasts were excluded. Non-leukemic cases consisted of those with less than 20% blasts and no prior history of acute leukemia.

### Cytogenetic and molecular genetic testing

Among 568 AML cases, 478 had concurrent cytogenetics available from karyotyping and/or fluorescence *in situ* hybridization analysis, and 476 had molecular genetic information available from Rapid Heme Panel testing at Brigham and Women’s Hospital^14^. **Supplementary Tables 3 and 4** provide the number of AML cases with different cytogenetic aberrancies and pathogenic variants, respectively. Cytogenetic aberrancies and pathogenic variants with a dataset prevalence of less than 2% were excluded from analysis due to lack of adequate sample size.

### Flow cytometry data preprocessing

FCS files outputted by BD FACSCanto™ II flow cytometers were loaded into Python scripts using the FlowKit package^15^ and normalized to marker mean values of 0 and standard deviation values of 1. No further preprocessing steps, including compensation, gating, or doublet exclusion, were performed.

### Encoder network development and pre-training

Two encoder neural networks were developed to analyze flow cytometry data from each of two tubes. These networks were designed as multilayer perceptrons (MLPs) with embedding modules based on periodic activation functions, which have been shown to perform similarly on tabular data to transformer models with significantly lower computational cost^16^. Specifically, MLP models with embedding function ReLU ◦ Linear ◦ Periodic (MLP-PLR) were used, with the Periodic function being:

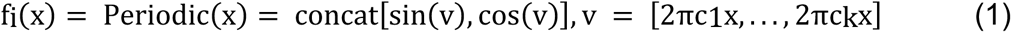

where *c_i_*are trainable parameters initialized from N(0,σ), and both σ and k are hyperparameters.

Encoder networks were pre-trained on all events from the respective tube of all 1,820 flow cytometry cases using Self-Supervised Contrastive Learning using Random Feature Corruption (SCARF)^17^. During pre-training, encoder hyperparameter optimization was performed using a random 1% subset of pre-training data for 100 iterations; **Supplementary Table 5** provides the hyperparameter search space and optimal values. Encoder networks with optimal hyperparameter values were then pre-trained on 100% of pre-training data.

### Predictive model development

Models for the prediction of presence/absence of acute leukemia, AML vs ALL, and presence/absence of cytogenetic aberrancies and pathogenic variants were designed as attention-based multi-instance learning models (ABMILMs) with a gated attention mechanism, as described by Ilse et al^12^. For each application, separate ABMILMs were developed for analysis of flow cytometry data from each tube; the outputs of each ABMILM were concatenated and inputted into a final MLP neural network to provide a class-level prediction. Hyperparameter optimization was performed for 50 iterations; **Supplementary Table 6** provides the hyperparameter search space. A negative log-likelihood loss function was used with class weights relative to the proportion of samples in each class. 5-fold cross validation of randomly-split training and validation datasets was performed for performance assessment of each model.

Due to memory constraints, a maximum threshold on the number of events per tube of 150,000 was implemented. For tubes with more than 150,000 events (impacting 151/1820 (8.3%) of tube 1 datasets and 152/1820 (8.4%) of tube 2 datasets), a random subset of 150,000 events was chosen.

### Analysis of model predictions

Machine learning models output the percentage probability of a particular sample being within each respective class. Model performance on validation datasets was assessed by calculating the area under the receiver operating characteristic curve (AUROC), as well as the accuracy, sensitivity, and specificity using a probability threshold which maximizes the Youden’s J statistic of the model.

### Analysis of model attention values and predictive power scores

Attention values from the two tube ABMILMs were analyzed to identify the relative importance of each flow cytometry event towards a model’s prediction for a particular sample. Attention values from each tube were normalized to range from (0,1) to improve visualization. Visualization of attention scores was achieved by coloring flow cytometry events based on normalized scores.

Predictive power scores (PPS), which are non-parametric scores detecting non-linear relationships between two variables, were calculated to assess the relationship between normalized attention weights and flow cytometry marker expression for individual cases^18^. PPS values rely on quantitative flow cytometric marker expression levels rather than arbitrary qualitative “positive/negative” gating of events. For a given model and flow cytometry marker, both the percentage of cases with non-zero PPS values, as well as the mean PPS value among non-zero cases, were used to assess the relative diagnostic importance of the marker to that model.

### Software and hardware

Python 3.8.10 and PyTorch 2.0.1+cu117 were used to develop and train neural network models^19^. All experiments were run on a Linux server with Intel Xeon Gold 6132 processor, 126 GB RAM, and 4 Tesla V100-PCIE-32GB GPUs.

## RESULTS

### Design of machine learning models for sample-level diagnosis and molecular characterization

We developed attention-based multi-instance learning models (ABMILMs) for each application of interest: (1) the prediction of presence *versus* absence of acute leukemia, (2) the differentiation of acute myeloid leukemia (AML) *versus* acute lymphoblastic leukemia (ALL), and (3) the prediction of presence *versus* absence of individual cytogenetic aberrancies and pathogenic variants (**Figure 1A**). These models are designed to input two-tube flow cytometry data from an individual sample, subsequently outputting the percentage probability of a sample being represented by each class (e.g., percentage probability that a case does *versus* does not represent acute leukemia). Data from each tube is first analyzed by an encoder, a neural network that identifies new features from the flow cytometry data for improved predictive capabilities. These new features are then analyzed by an attention module, which decides which individual flow cytometry events are important *versus* not important for the diagnosis, and accordingly assigns each event a weight based on this perceived importance. Using these weights along with the encoder-outputted features, pooled features from each tube are obtained. The pooled features from both tubes are concatenated and finally analyzed by an integrating neural network to provide the final prediction for the entire sample.

**Figure 1.**
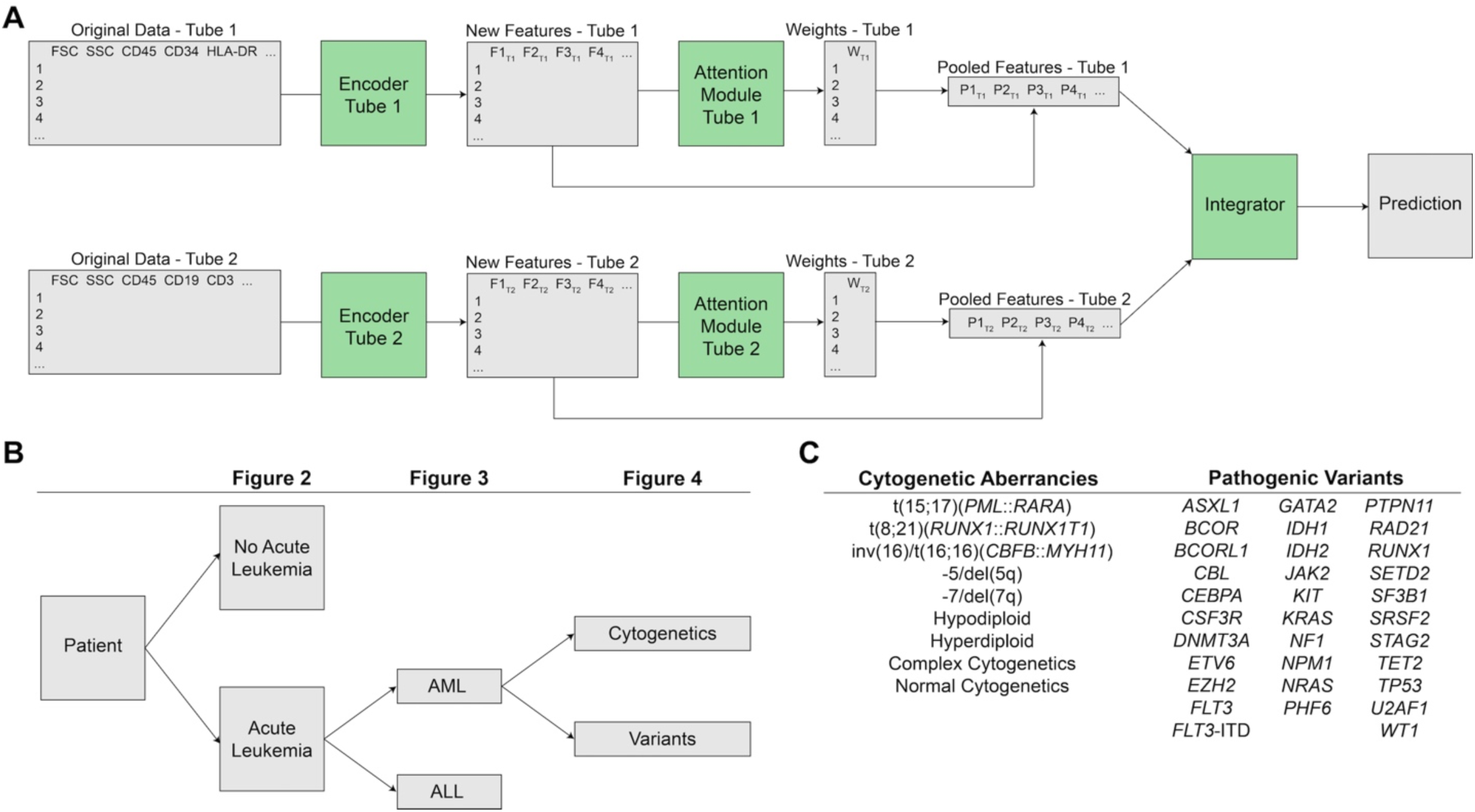
Workflow for automated machine learning-based diagnosis and molecular characterization of acute myeloid leukemia from flow cytometry data. (A) Schematic of attention-based multi-instance learning models. Flow cytometry data extracted from FCS files from two-tube leukemia panels are provided as input to encoder neural networks which extract new features from the data in each respective tube. These new features are analyzed by attention modules which designate a weight for each flow cytometry event based on the perceived importance of the event in overall case diagnosis by the model. These weights, along with the new features, are used to extract pooled features from each tube. Pooled features from the two tubes are combined and analyzed by a final neural network which provides a sample-level prediction for the sample. **(B)** Workflow for sample diagnosis and molecular characterization. The first model predicts the presence or absence of acute leukemia. Acute leukemia cases are then analyzed by the next model to differentiate between AML and ALL cases. Finally, AML cases are analyzed by a set of individual models that predict the presence or absence of cytogenetic aberrancies and pathogenic variants. **(C)** List of cytogenetic aberrancies and pathogenic variants tested for in AML cases. Only variants with dataset prevalence of at least 2% were chosen to ensure adequate sample size.

A stepwise workflow for the diagnosis and molecular characterization of individual cases was developed, first identifying which cases are positive for acute leukemia, which of those represent AML cases, and finally which cytogenetic aberrancies and pathogenic variants each AML case has (**Figure 1B**). **Figure 1C** shows the specific cytogenetic aberrancies and pathogenic variants for which predictive models were developed. Only variants with dataset prevalence of at least 2% were chosen to ensure adequate sample size.

### Machine learning model for the diagnosis of acute leukemia

A machine learning model was first developed to distinguish between non-leukemic cases and cases with acute leukemia. This model demonstrated strong predictive performance overall (AUROC 0.961 ± 0.011, accuracy 0.903 ± 0.013), as well as individually on leukemic (sensitivity 0.881 ± 0.020 at Youden’s J) and non-leukemic (specificity 0.925 ± 0.027 at Youden’s J) cases (**Figure 2A**). **Figure 2A** also shows that by adjusting the predictive probability threshold along the ROC curve, model sensitivity can be significantly increased with a relative tradeoff in specificity; this could allow the model to be used in more of a triaging/screening role^20^. Analysis of flow cytometry event attention values for individual leukemia cases shows that, analogous to manual diagnosis by a hematopathologist, the model learns to pay most attention to blast cells for diagnosis, with mixed attention to monocytic cells, and less attention to mature lymphocytes and granulocytes (**Figure 2B**).

**Figure 2.**
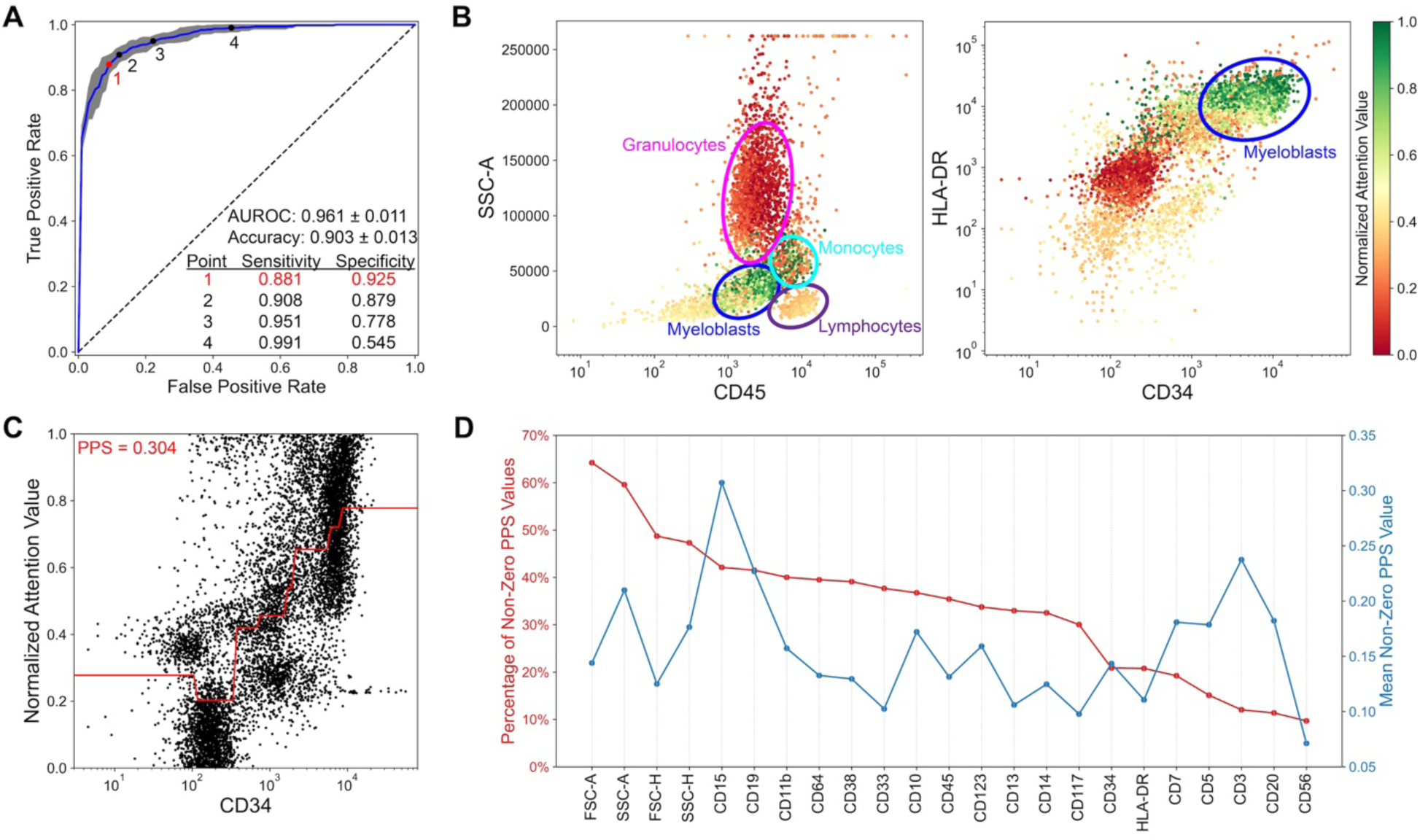
Machine learning model for the diagnosis of acute leukemia. (A) Receiver operating characteristic (ROC) curve for the model predicting the presence or absence of acute leukemia from flow cytometry samples. Blue line represents the mean of five training/validation data splits, and gray outline represents the standard error. Model AUROC and accuracy are provided. Numbered points along the ROC curve indicate varying sensitivity and specificity values with different predictive probability threshold values. Point #1 (red) indicates the point along the ROC curve which maximizes Youden’s J, whereas points #2-4 (black) indicate those with sensitivity values of 90%, 95%, and 99%, respectively. **(B)** Analysis of model attention values for a representative AML case, which was correctly identified by the model as being positive for acute leukemia. Plots of side scatter (SSC-A) *versus* CD45, and HLA-DR *versus* CD34, are provided. Individual events are colored based on the normalized attention value assigned by the model to each flow cytometry event. Cell populations of interest are illustrated. **(C)** Diagram demonstrating an example calculation of predictive power scores (PPS) for quantifying the relationship between attention values and flow cytometry marker expression for an individual case. Red line represents the non-linear relationship identified between the two variables. The PPS score that quantifies this relationship on a scale from 0-1 is provided. **(D)** Diagram summarizing the PPS values for all flow cytometry markers in the prediction of the presence or absence of acute leukemia. Values show (red, left) the percentage of non-zero PPS values across all samples, and (blue, right) the mean non-zero PPS value across all samples. Markers are ordered based on descending value of the percentage of non-zero PPS values.

To assess the overall importance of each marker towards model predictions, predictive power scores (PPS) were calculated to quantify the relationship between attention values and flow cytometry marker expression for individual cases; by fitting a non-parametric model to the data which can capture non-linear relationships, this approach provides a numerical score analogous to a correlation coefficient which represents the strength of the relationship between attention values and marker expression (**Figure 2C**). PPS values across cases were averaged by assessing both the percentage of non-zero PPS values per flow cytometry marker, as well as the mean non-zero PPS value for each marker (**Figure 2D**).

PPS value analysis for this model shows that the machine learning model utilized forward scatter (FSC) and side scatter (SSC) information in a larger percentage of cases than any other flow cytometry markers. Among cell surface markers, CD15 (sialyl Lewis x), a marker for human myeloid cells, was associated with both the highest percentage of non-zero PPS values, as well as the highest magnitude PPS values^21^. CD19, critical for the diagnosis of B-ALL and AML with t(8;21)(*RUNX1*::*RUNX1T1*), also showed strong predictive performance^22^. Interestingly, flow cytometry markers often used for manual blast identification in AML, including CD45, CD34, and HLA-DR, were among the least important cell surface markers used by the model^23^. Finally, while CD3 was among the least used markers by the model, it displayed the second highest mean value among non-zero cases; this likely represents the use of CD3 in identification of the limited number of T-ALL cases in the dataset.

### Machine learning model for classification of AML *versus* ALL cases

Performance of a machine learning model for classification of AML *versus* ALL (including B-ALL and T-ALL) showed equally strong performance (AUROC 0.965 ± 0.015, accuracy 0.922 ± 0.025) as initial diagnosis of acute leukemia (**Figure 3A**). While events representing blast cells still had comparably high attention values for most cases, other cell types appeared to provide similar or larger importance to model predictions, including mature granulocytes with high SSC values, as well as a subset of mature lymphocytes (**Figure 3B**). This suggests that the immunophenotypic profile of cell types other than blast cells may provide an underexplored role in the diagnosis of acute leukemia cases.

**Figure 3.**
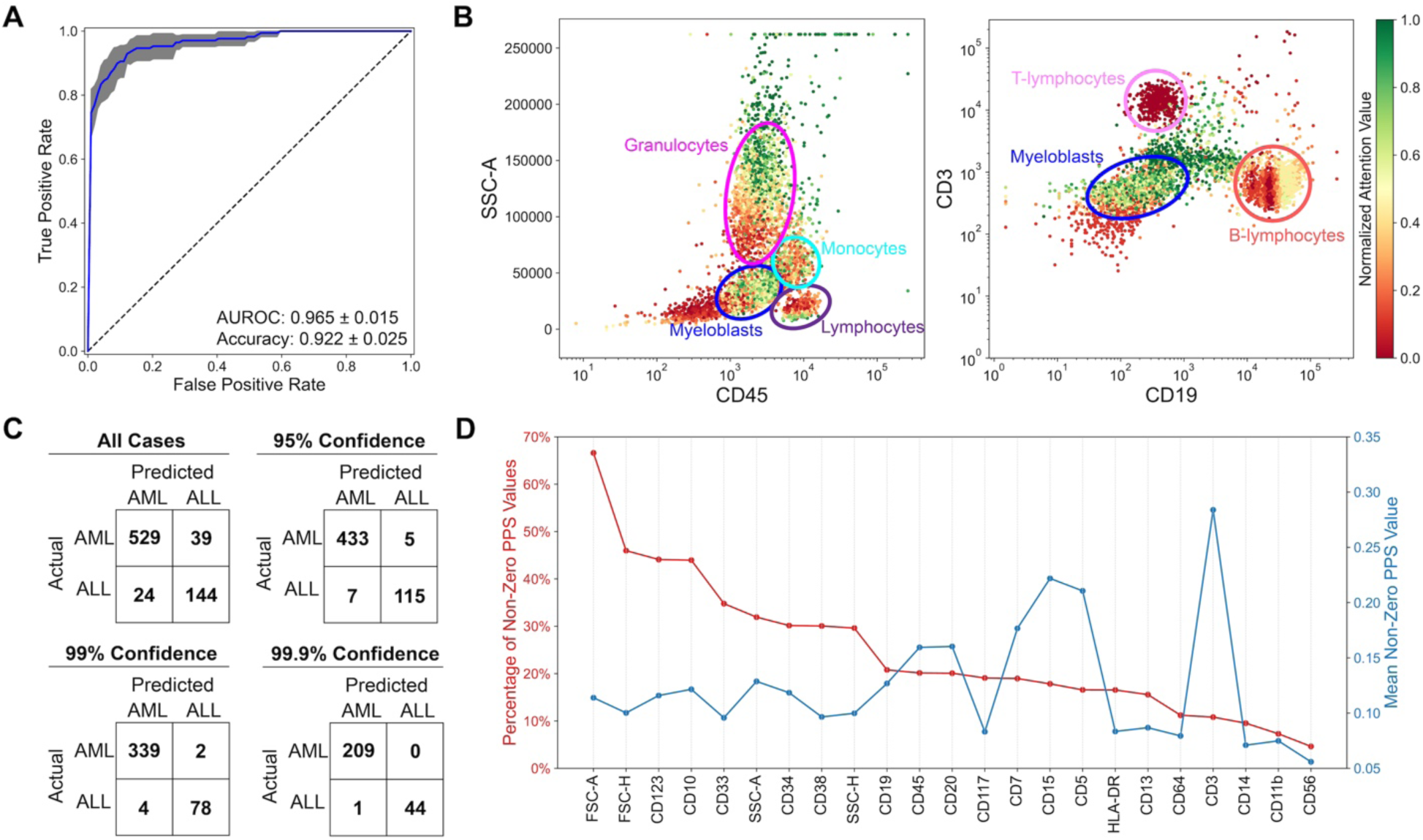
Machine learning model for the classification of AML *versus* ALL. (A) Receiver operating characteristic (ROC) curve for the model predicting the presence of AML *versus* ALL from flow cytometry samples. Blue line represents the mean of five training/validation data splits, and gray outline represents the standard error. AUROC and accuracy values for the model are provided. **(B)** Analysis of model attention values for a representative AML case, which was correctly identified by the model as being positive for AML. Plots of side scatter (SSC-A) *versus* CD45, and CD3 *versus* CD19, are provided. Individual events are colored based on the normalized attention value assigned by the model to each flow cytometry event. Cell populations of interest are illustrated. **(C)** Confusion matrices showing the accuracy of model predictions at different model confidence thresholds. Cases where the predicted probability of a case being either an AML or ALL case is above the provided threshold are included. Rows represent the actual class and columns represent the predicted class. **(D)** Diagram summarizing the PPS values for all flow cytometry markers in the prediction of the presence of AML *versus* ALL. Values show (red, left) the percentage of non-zero PPS values across all samples, and (blue, right) the mean non-zero PPS value across all samples. Markers are ordered based on descending value of the percentage of non- zero PPS values.

Despite the model’s strong accuracy, misclassifications were still made for cases with lower model confidence. An analysis was performed to determine at what model- outputted confidence level the machine learning model can make highly accurate predictions for classification of AML *versus* ALL (**Figure 3C**). For cases with model confidence of at least 95% (560/736, 76.1% of total cases), the model’s cumulative accuracy across all validation sets increased from 91.4% to 97.9%. Furthermore, at a confidence threshold of 99.9% (254/736, 34.5% of total cases), only one misclassification was made, with a model accuracy of 99.6%. Thus, while employing such a high confidence threshold may limit model generalizability, it may be favored to make highly accurate predictions in particularly challenging cases.

Analysis of model PPS values shows that FSC values carried small but non-zero importance in most cases, representative of the fact that myeloblasts and lymphoblasts often exhibit subtle differences in cell size^24^ (**Figure 3D**); interestingly, side scatter showed lower relative importance despite differences in granularity between myeloblasts and lymphoblasts. CD123 (IL3 receptor), which shows expression in most AML and B-ALL cases but not T-ALL cases, as well as CD10, which is commonly expressed in ALL but not AML, yielded relatively high importance values^25,26^. Finally, CD15 as well as T-cell markers (CD3, CD5, and CD7), demonstrated high importance in a small subset of AML and T-ALL cases.

### Prediction of cytogenetic aberrancies and pathogenic variants in AML cases

Individual machine learning models predicting the presence or absence of 9 cytogenetic aberrancies and 32 pathogenic variants among AML cases were developed (**Figure 4A**). The model for prediction of t(15;17)(*PML*::*RARA*), which has a distinct immunophenotypic profile with negative CD34 in the hypergranular variant, negative HLA-DR, and positive CD117 expression, showed the strongest performance among all variants tested (AUROC 0.929 ± 0.032, accuracy 0.885 ± 0.071)^27^; the model performed well for both positive (sensitivity 0.925 ± 0.061) and negative (specificity 0.882 ± 0.083) cases (**Figure 4B**). Other models with strong performance are associated with variants with known immunophenotypic signatures, including t(8;21)(*RUNX1*::*RUNX1T1*) (AUROC 0.814 ± 0.050) which is associated with CD19 and CD56 expression^28^, and *NPM1* variants (AUROC 0.807 ± 0.020) which are associated with CD19 and CD4 expression as well as monocytic markers^29^. 9/41 (22.0%) of variant models demonstrated an AUROC > 0.7, and 32/41 (78.0%) of variant models demonstrated an AUROC > 0.6.

**Figure 4.**
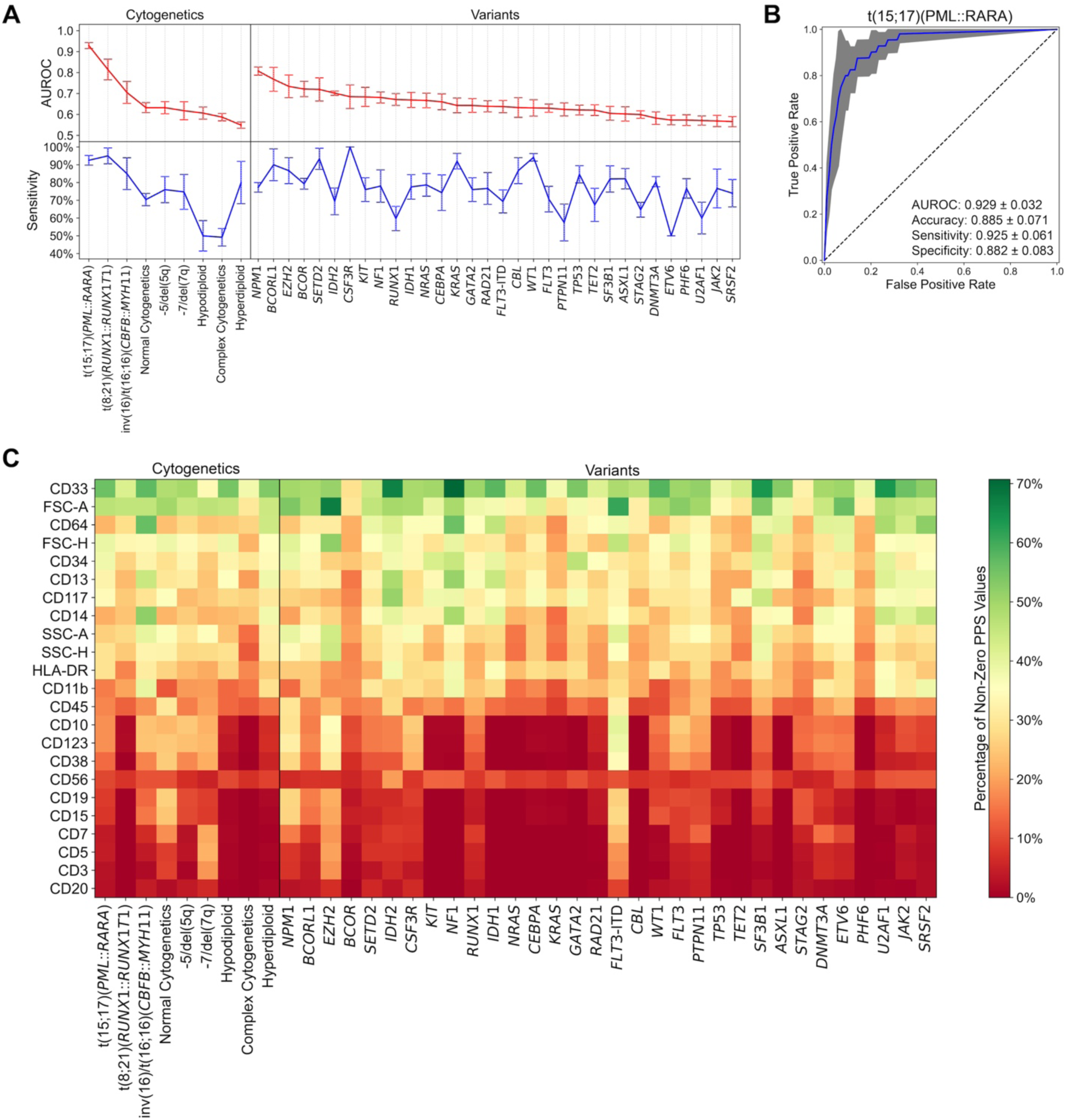
Machine learning models for predicting the presence or absence of cytogenetic aberrancies and pathogenic variants in AML cases. (A) Performance metrics (AUROC: red, top; sensitivity: blue, bottom) of individual machine learning models predicting the presence or absence of a particular cytogenetic aberrancy or pathogenic variant. Bar graph represents the mean ± one standard error. Cytogenetic aberrancies and pathogenic variants are ordered based on descending AUROC value. **(B)** Receiver operating characteristic (ROC) curve for the model predicting the presence or absence of the t(15;17)(*PML*::*RARA*) translocation from flow cytometry samples. Blue line represents the mean of five training/validation data splits, and gray outline represents the standard error. AUROC, accuracy, sensitivity, and specificity values for the model are provided. **(C)** Heatmap showing the percentage of non-zero PPS values representing the importance of each flow cytometry marker for all predictive models. Flow cytometry markers are ordered (top to bottom) based on decreasing mean value of the percentage of non-zero PPS values among all models.

PPS score analysis across all predictive models shows that flow cytometry markers can be placed into three groups based on relative importance for model predictions (**Figure 4C**). A select few markers, including forward scatter (FSC-A/H), and CD33, demonstrated strong predictive utility in almost all models. While CD33 expression has been implicated in *NPM1*- and *FLT3*-mutated AML^30,31^, these results appear to associate CD33 with many other cytogenetic and mutational variants, with implications for targeted therapy^32^. Many other markers, including B-cell (CD19 and CD20) and T-cell (CD3, CD5, and CD7) markers, exhibit very little predictive value for most models. Finally, some markers show strong importance for individual models but otherwise weak importance. For example, monocytic markers CD14 and CD64 had high importance scores for prediction of inv(16)/t(16;16)(*CBFB*::*MYH11*), which often exhibits monocytic differentiation^33^. Additionally, CD117 had a relatively high importance score for prediction of *IDH2* mutations; prior studies have associated *IDH2* R172 mutations with high CD117 expression^34^, and *IDH2*-targeting therapies have been shown to decrease CD117 expression in AML^35^.

### Visualization and analysis of model outputs for individual patient cases

Figure 5 provides an example of a model-generated report for a representative case of acute promyelocytic leukemia with confirmed presence of t(15;17)(*PML*::*RARA*) by cytogenetics/FISH, but which had significant diagnostic uncertainty during manual interpretation. The model correctly identified the presence of leukemia (predicted probability p = 98.6%) and AML rather than ALL (p = 100.0%). The model also correctly predicted the presence of the t(15;17)(*PML*::*RARA*) translocation (p = 75.4%), as well as a *FLT3* pathogenic variant (albeit with low predictive confidence; p = 50.4%).

**Figure 5.**
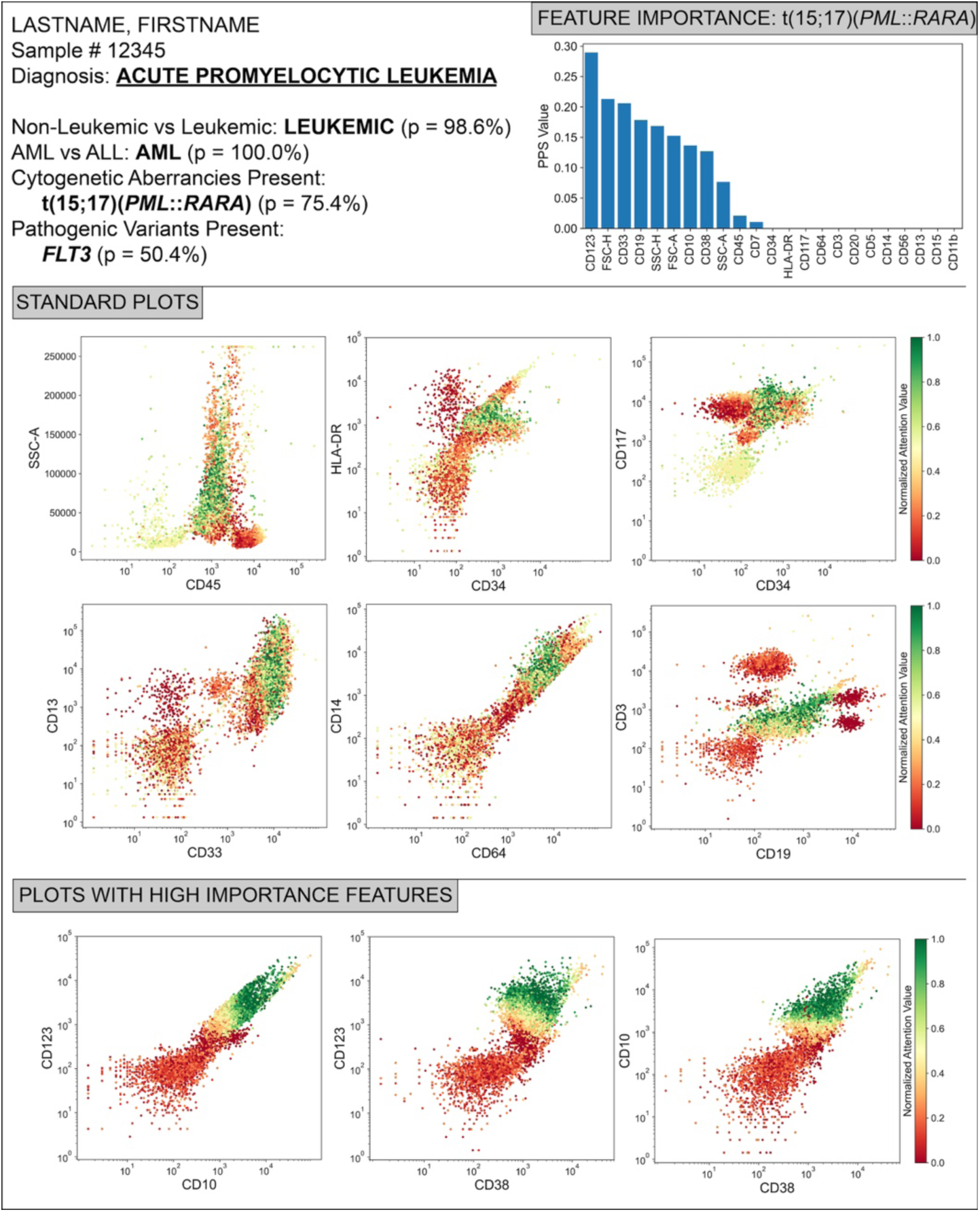
Example model-generated report of representative acute promyelocytic leukemia case. (Top-Left) The final diagnosis of acute promyelocytic leukemia, as well as model predictions and associated confidence values along the entire computational pipeline, are shown. **(Top-Right)** PPS values for flow cytometry markers in the prediction of t(15;17)(*PML*::*RARA*) for this case. **(Bottom)** Analysis of model attention values and expression of flow cytometry markers conventionally used in diagnosis of acute promyelocytic leukemia, as well as those identified as diagnostically important for this particular case.

To illustrate the utility of this computational pipeline for identification of cytogenetic aberrancies/pathogenic variants in challenging cases, flow cytometry marker expression and corresponding event attention values were further analyzed. Significant model attention was placed on the population of CD45-dim immature cells that were characterized as SSC(increased)/CD45(dim)/CD34(variable)/HLA-DR-/CD117+/CD13(variable)/CD33+/CD14-/CD64+/CD3-/CD19-. Although most of this immunophenotype is consistent with hypergranular APML, the variable CD34 positivity is quite infrequent in hypergranular APML (only approximately 16% of cases)^23^ and likely led to the diagnostic uncertainty about the presence of t(15;17)(*PML*::*RARA*).

PPS value analysis showed that many flow cytometry markers not commonly utilized in assessment of APML, including CD123, CD10, and CD38, were associated with high feature importance for diagnosis of this case. Figure 5 shows that the population of promyelocytes demonstrates high expression of these markers, and the model’s attention values are highly correlated with marker expression. Additionally, conventionally used markers for APML including CD45, CD34, and HLA-DR surprisingly carried small or no importance in the case diagnosis. Thus, the machine learning model was able to correctly predict the presence of t(15;17)(*PML*::*RARA*) by developing a previously-uncharacterized association between this translocation and increased expression of CD123, CD10, and CD38, rather than relying on markers more commonly used during manual analysis.

## DISCUSSION

While statistical and machine learning approaches have been extensively applied to analysis of flow and mass cytometry data^6^, these prior approaches carry significant shortcomings. Dimensionality reduction methods for data visualization (including PCA, tSNE, and UMAP), as well as unsupervised methods for cell population clustering (including flowSOM^36^, FLOCK^37^, and PhenoGraph^38^) are capable of identifying clinically-relevant cell populations, but are not primarily designed for classification applications such as predicting the presence or absence of a disease state or cytogenetic/molecular aberrancies. Past supervised learning methods including linear discriminant analysis^39^ and support vector machines^40^ often underperform current state-of-the-art deep learning methods in classification tasks. Most deep learning approaches to flow and mass cytometry data, including CellCNN^41^ and Deep CNN^8^, have utilized convolutional neural networks, a machine learning architecture that is well suited for image classification tasks, but is not designed for analysis of tabular data such as flow cytometric data^42^. Deep learning approaches specifically designed for tabular data have yet to be applied for the diagnosis and molecular characterization of hematolymphoid malignancies from flow cytometry data. Additionally, attention-based multi-instance learning, a deep learning approach which learns to identify (or pay “attention” to) particular data components (e.g., individual flow cytometric events within a sample) which carry the most diagnostic value in sample classification, has yet to be applied to analysis of flow cytometry data.

This study is, to our knowledge, the first to apply attention-based multi-instance learning to flow cytometry data for automated hematopathology diagnostics (Figure 1). We demonstrated that this approach provides highly accurate predictions on the presence/absence of acute leukemia (Figure 2), as well as for discriminating between AML and ALL (Figure 3). We additionally show the feasibility of using these models trained solely on flow cytometry data to provide insight on the presence or absence of clinically relevant cytogenetic aberrancies and pathogenic variants in AML-positive samples (Figure 4). Finally, we provide a representative example of how model outputs can be visualized and incorporated into a hematopathologist’s workflow for improved patient diagnosis, and for identification of novel associations between flow cytometry marker expression and particular molecular aberrancies (Figure 5).

This automated approach to flow cytometry-based diagnosis and molecular characterization of AML yields several potential use cases. For clinicians, the computational pipeline provides accurate and rapid predictions (within minutes after flow cytometry data collection) on disease status, including insight into the molecular subtype, of their patients, allowing for faster initiation of optimal treatment protocols. Additionally, model output visualizations can be tailored for clinicians to provide succinct yet insightful information into patients’ immunophenotype with therapeutic implications. For hematopathologists, the pipeline can be used as a triaging tool to rule out cases with very low predicted probabilities of acute leukemia, as well as a diagnosis assistance tool providing unbiased predictions of disease state with associated data visualizations for data analysis. For hematology laboratories as well as low-resource settings, the pipeline provides a software- and personnel-agnostic approach to flow cytometry-based diagnosis without need for manual intervention. Finally, for biomedical researchers, this tool provides information about novel associations between myeloblast cell surface marker expression and AML molecular subtypes which could be further explored as therapeutic targets.

Future efforts will continue to improve the generalizability of this approach toward particular technical considerations, use cases, and application areas. Currently, our computational pipeline does not discriminate between new *versus* recurrent cases of AML, and our model for predicting the presence or absence of acute leukemia requires a 20% blast count for positive cases. Modifications to model architecture including utilizing the patient’s previously characterized immunophenotype could be made to improve performance on recurrent cases, as well as for minimal/measurable residual disease assessment of minute cell populations. Additionally, these models were trained on a set flow cytometry panel with no changes in markers, antibodies, or fluorochromes. Incorporation of additional model elements which learn associations between flow cytometry markers using cross-institutional data could be a potential approach to allow for variance in flow cytometry panel construction, the number of colors per tube, and the use of multiple markers per channel. Lastly, addition of CNNs into the model architecture would allow for integration of flow cytometry data with cell morphologic data from blood and bone marrow aspirate smears, which could provide improved diagnostic accuracy.

Overall, this study represents a significant advancement in digital hematopathology by demonstrating the feasibility of using deep learning toward automated diagnosis and molecular characterization of AML cases solely using flow cytometry data. This automated approach provides many advantages and potential use cases compared to manual flow cytometric assessment for clinicians, hematopathologists, and the hematology laboratory. Continued improvements and integration with other data modalities including cell morphologic assessment will further enhance model performance and generalizability, ultimately moving these automated approaches closer to clinical use.

## ACKNOWLEDGEMENTS

This study was supported by National Institutes of Health National Cancer Institute grants U01CA220401 and U24CA19436201.

## ETHICS APPROVAL AND CONSENT TO PARTICIPATE

This study was approved by the Institutional Review Board. The study was performed in accordance with the Declaration of Helsinki.

## AUTHORSHIP CONTRIBUTIONS

J.E.L. and O.P. conceived the study; J.E.L. collected data and developed software; L.A.D.C. provided computational resources; J.E.L., L.A.D.C., D.L.J., and O.P. analyzed and interpreted the results; J.E.L. and O.P. wrote the manuscript. All authors read and approved the final paper.

## DATA SHARING STATEMENT

Data sharing requests should be sent to Olga Pozdnyakova.

---

**Supplementary Table 1.** Flow cytometry parameters and markers used in two-tube acute leukemia panel from this study.

| **Tube 1** |  | **Tube 2** |  |
| --- | --- | --- | --- |
| *Marker* | *Fluorophore* | *Marker* | *Fluorophore* |
| FSC-A | - | FSC-A | - |
| FSC-H | - | FSC-H | - |
| SSC-A | - | SSC-A | - |
| SSC-H | - | SSC-H | - |
| CD45 | V500-C | CD45 | V500-C |
| CD34 | FITC | CD3 | BV421 |
| HLA-DR | PerCP-Cy5.5 | CD5 | PE-Cy7 |
| CD11b | APC | CD7 | BV605 |
| CD13 | PE-Cy7 | CD10 | APC-R700 |
| CD14 | APC-H7 | CD15 | FITC |
| CD33 | PE | CD19 | PE |
| CD56 | BV421 | CD20 | APC-H7 |
| CD64 | APC-R700 | CD38 | PerCP-Cy5.5 |
| CD117 | BV605 | CD123 | APC |

**Supplementary Table 2.** Number of patients in respective diagnostic categories from this study.

| **Category** | **# Cases** | **% Cases** |
| --- | --- | --- |
| All cases (used for encoder pre-training) | 1820 |  |
| Cases used for model training | 1468 |  |
| No acute leukemia | 732 | 49.9% |
| Acute leukemia | 736 | 50.1% |
| ALL | 168 | 22.8% |
| AML | 568 | 77.2% |
| Cytogenetic aberrancy data available | 478 | 84.2% |
| Pathogenic variant data available | 476 | 83.8% |

**Supplementary Table 3.** Number of cases based on the presence of cytogenetic aberrancies.

| **Cytogenetic Aberrancy** | **# Cases** | **% Cases** |
| --- | --- | --- |
| Normal Karyotype | 159 | 33.3% |
| Complex Karyotype | 104 | 21.8% |
| -5/del(5q) | 46 | 9.6% |
| -7/del(7q) | 42 | 8.8% |
| t(15;17)(*PML::RARA*) | 40 | 8.4% |
| Hypodiploid Karyotype | 34 | 7.1% |
| t(8;21)(*RUNX1::RUNX1T1*) | 22 | 4.6% |
| inv(16)/t(16;16)(*CBFB::MYH11*) | 19 | 4.0% |
| Hyperdiploid Karyotype | 15 | 3.1% |

**Supplementary Table 4.** Number of cases based on presence of pathogenic variants.

| **Variant** | **# Cases** | **% Cases** |
| --- | --- | --- |
| *FLT3-ITD* | 131 | 27.5 |
| *NPM1* | 123 | 25.8 |
| *DNMT3A* | 98 | 20.6 |
| *TET2* | 97 | 20.4 |
| *RUNX1* | 87 | 18.3 |
| *FLT3* | 75 | 15.8 |
| *NRAS* | 75 | 15.8 |
| *IDH1* | 72 | 15.1 |
| *TP53* | 65 | 13.7 |
| *SRSF2* | 61 | 12.8 |
| *IDH2* | 58 | 12.2 |
| *ASXL1* | 57 | 12.0 |
| *WT1* | 53 | 11.1 |
| *PTPN11* | 39 | 8.2 |
| *BCOR* | 39 | 8.2 |
| *CEBPA* | 31 | 6.5 |
| *SF3B1* | 28 | 5.9 |
| *STAG2* | 26 | 5.5 |
| *KRAS* | 26 | 5.5 |
| *GATA2* | 25 | 5.3 |
| *KIT* | 24 | 5.0 |
| *NF1* | 23 | 4.8 |
| *U2AF1* | 20 | 4.2 |
| *JAK2* | 19 | 4.0 |
| *PHF6* | 17 | 3.6 |
| *EZH2* | 15 | 3.2 |
| *CBL* | 13 | 2.7 |
| *SETD2* | 13 | 2.7 |
| *RAD21* | 12 | 2.5 |
| *ETV6* | 10 | 2.1 |
| *BCORL1* | 10 | 2.1 |
| *CSF3R* | 10 | 2.1 |

**Supplementary Table 5.** Hyperparameter values from encoder pre-training.

| Hyperparameter | Search Range | Optimal Value – Tube 1 | Optimal Value – Tube 2 |
| --- | --- | --- | --- |
| k (periodic activation function, # features / 2) | Int(1,128) | 50 | 102 |
| $\sigma$ (periodic activation function, $\max(c_i)$) | Float(0.01,100), log range | 17.875532 | 0.344150 |
| # neurons in linear layer of PLR embedding | $e^{\text{Int}(0,7)}$ | $e^4$ | $e^6$ |
| # MLP blocks | Int(1,16) | 1 | 7 |
| # neurons in linear layer of MLP block | $e^{\text{Int}(0,10)}$ | $e^6$ | $e^9$ |
| Dropout rate of MLP block | Float(0,0.5) | 0.425532 | 0.038182 |
| Learning Rate | Float($5e^{-5}$, $5e^{-3}$) | $1.824962e^{-4}$ | $8.712371e^{-5}$ |
| Weight decay | Float($1e^{-6}$, $1e^{-3}$) | $2.252650e^{-5}$ | $4.066816e^{-5}$ |
| Corruption rate | Float(0.2,0.8) | 0.210043 | 0.201649 |

**Supplementary Table 6.** Hyperparameter values from multi-instance learning model training.

| Hyperparameter | Search Range |
| --- | --- |
| # hidden layers, attention module, tube 1 | Int(0,10) |
| # hidden layers, attention module, tube 2 | Int(0,10) |
| # layers, integrator MLP | Int(1,16) |
| # neurons per layer, integrator MLP | $e^{\text{Int}(0,10)}$ |
| Dropout rate of MLP block | Float(0,0.5) |
| Learning Rate | Float($5e^{-5}$, $5e^{-3}$) |
| Weight decay | Float($1e^{-6}$, $1e^{-3}$) |

